# Latent generative modeling of long genetic sequences with GANs

**DOI:** 10.1101/2024.08.07.607012

**Authors:** Antoine Szatkownik, Cyril Furtlehner, Guillaume Charpiat, Burak Yelmen, Flora Jay

**Author notes:** Corresponding author: Antoine Szatkownik < >. These authors contributed equally (Burak Yelmen, Flora Jay).

## Abstract

Synthetic data generation via generative modeling has recently become a prominent research field in genomics, with applications ranging from functional sequence design to high-quality, privacy-preserving artificial in silico genomes. Following a body of work on Artificial Genomes (AGs) created via various generative models trained with raw genomic input, we propose a conceptually different approach to address the issues of scalability and complexity of genomic data generation in very high dimensions. Our method combines dimensionality reduction, achieved by Principal Component Analysis (PCA), and a Generative Adversarial Network (GAN) learning in this reduced space. Using this framework, we generated genomic proxy datasets for very diverse human populations around the world. We compared the quality of AGs generated by our approach with AGs generated by the established models and report improvements in capturing population structure, linkage disequilibrium, and metrics related to privacy leakage. Furthermore, we developed a frugal model with orders of magnitude fewer parameters and comparable performance to larger models. For quality assessment, we also implemented a new evaluation metric based on information theory to measure local haplotypic diversity, showing that generative models yield higher diversity than real genomes. In addition, we addressed the shrinkage issue associated with PCA and generative modeling, examined its relation to the nearest neighbor resemblance metric, and proposed a resolution. Finally, we evaluated the effect of different binarization methods on the quality of the output AGs.

## 1 Introduction

Growing advances in DNA sequencing have spurred a data surge, elevating machine learning’s importance in genomics (Yelmen and Jay, 2023; Korfmann, 2023; Huang et al., 2023). However, a substantial amount of these data, held by companies, institutions or governmental agencies, are tied to privacy concerns impeding their accessibility. In contemporary populations genomics, the unsupervised training of generative neural networks on genomes mainly aims at enhancing visualization by revealing fine-scale population structure (Battey et al., 2021; Meisner and Albrechtsen, 2022; Geleta et al., 2023), inferring evolutionary histories (Wang et al., 2021; Riley et al., 2024), or generating realistic synthetic data (Yelmen et al., 2021; Perera et al., 2022; Booker et al., 2023; Yelmen et al., 2023; Dang et al., 2023; Zhang et al., 2024; Burnard et al., 2023; Ahronoviz and Gronau, 2024; Szatkownik et al., 2024). For the latter goal, an alternative paradigm consists of simulations based on the resampling of known haplotypes (Su et al., 2011; Tang and Liu, 2019; Dimitromanolakis et al., 2019; Wharrie et al., 2023). Generated synthetic genomes, or artificial genomes (AGs), can potentially become privacy-preserving alternatives to real biobanks and be utilized for genome-wide association studies (GWAS), local ancestry inference, genomic imputation or natural selection scans (Yelmen et al., 2021).

Generative neural networks are a class of deep-learning based algorithms, that once trained, can be used to map noise to a data distribution that should resemble as closely as possible to the real training data distribution. Several models have been developed over the years with varying degrees of success, ranging from variational autoencoders (VAEs) (Kingma and Welling, 2014), generative adversarial networks (GANs) (Goodfellow et al., 2014), Restricted Boltzmann Machines (RBMs) (Smolensky, 1986; Ackley et al., 1985) to diffusion models (Ho et al., 2020) and language models (Zhang et al., 2024). In this paper, we focus on GANs, which consist of a circuit of two neural networks competing with each other where one, the discriminator, is assigned the task of distinguishing fake from real data while the other one, the generator, is simulating real data with synthetic data via a generative process whose purpose is to trick the discriminator.

In this context of generative AI for genomics, Yelmen et al. (2023) tackled the challenging problem of generating very high dimensional genetic data despite small sample sizes by designing and training convolutional Generative Adversarial Networks (GANs), convolutional Variational Autoencoders (VAEs), and conditional Restricted Boltzmann Machines (RBMs). Unlike previous fully connected architectures (Yelmen et al., 2021), the proposed scheme of convolving along the genomic sequences combined with location-specific variables is computationally feasible (Ausmees and Nettelblad, 2022; Yelmen et al., 2023), yet it might still pose challenges when targeting dense and intrinsically complex whole genomes. These difficulties drive the need to seek alternative methods for large genome-wide frameworks.

The recent staggering success of generative AIs for high-resolution image synthesis such as *DALL-E* (Ramesh et al., 2021) or *Stable Diffusion* (Rombach et al., 2022) roots in a two-stage process: learning a good latent representation of the data and then training a generative model in the resulting lower-dimensional subspace. This paradigm sparked growing attention within the ML community, as illustrated by the *NeurIPS Tutorial* set up in 2023, *Latent Diffusion Models: Is the Generative AI Revolution Happening in Latent Space?* which attracted thousands of people (LDM, 2023). Inspired by this, we call *latent generative models*, the association of a compression step followed by generative modeling in that space. This framework enables high-quality synthetic data generation while bypassing massive volumes of computations. Although it has been suggested that these two steps could be learned jointly, results were less prominent (Vahdat et al., 2021; Rombach et al., 2022).

In addition, despite being straightforward architectures to implement, train and optimize, fully-connected (FC) architectures are not practical for direct application to large-scale genetic data. For instance, a simple model consisting of just one dense linear layer with *n* = 1000 hidden variables and dealing with genetic sequences of *L* single nucleotide polymorphisms (SNPs), with *L* potentially of the order of millions (e.g., *L* = 1M), would have *L × n* = 1B parameters. In a latent generative modeling framework, boiling the important information down to a smaller space bypasses this large computational overhead and renders FC affordable.

To address the high-dimensional generative task, we thus propose to project the genetic data into a lower-dimensional subspace through dimension reduction and to train a GAN within this reduced space. Principal Component Analysis (PCA) was preferred for its easy and faithful reconstruction from Principal Component (PC) space to data space, unlike t-SNE and UMAP, which either lack invertibility or result in poor reconstruction. While autoencoders are also of interest, they are heavily parameterized, leading to long training times, and require careful regularization adjustments. On the contrary, PCA is nonparametric, efficiently computed in minimal time, and widely used in population genetics studies since ancestry-linked genomic variation is well represented in PC space (Tian et al., 2008; Price et al., 2010; Lao et al., 2008; Novembre et al., 2008; Yu et al., 2008; Menozzi et al., 1978; Patterson et al., 2006; Hanotte et al., 2002; Price et al., 2006). In the principal subspace, each new dimension is a linear combination of the input features, therefore the notion of locality along features disappears when replacing the genomic sequences by their PC scores. This makes dense layers or attention layers ideal candidates for the generative network architecture. Importantly, although PCA is linear, the generator output is not, as the GAN can learn non-linearities. This preserves our capacity to capture the complex structure of genomic sequences.

In this study, we introduce three latent generative modeling methods, a PCA Wasserstein GAN with gradient penalty (PCA-WGAN), a chained alternative designed to preserve local information (Glocal-PCA-WGAN), and a frugal version with fewer parameters (Light PCA-WGAN). We showcase the usefulness of our approaches through comparative analyses with previous models on The 1000 Genomes Project Consortium (2010) genetic dataset encompassing diverse human populations and demonstrate improvements on several population genetics summary statistics and metrics related to privacy leakage.

## 2 Materials and Methods

### 2.1 Overview

Our general approach for generating realistic genomic sequences consists of (i) reducing the training dataset dimensions through principal component analysis; (ii) training a WGAN on these PC scores instead of the haplotypes; (iii) generating synthetic PC scores by sampling the generator; (iv) inverse-transforming these scores back to the genetic space, yielding synthetic genome samples; (v) assessing the quality of the artificial genomes using multiple descriptive population genetics statistics (**FIG. S1**).

### 2.2 Dataset description

The 1000 Genomes project is a database of human genome sequences obtained by sampling individuals from 26 populations spread worldwide, with a number of samples for each population ranging from one to roughly a hundred. The 1000 Genomes dataset used in this study was curated by Yelmen et al. (2023). It consists of 2504 individuals corresponding to 5008 phased haplotypes (*i.e*. for each individual there is one reconstructed haplotype coming from the mother and one from the father), and contains 65,535 contiguous SNPs spanning chr1:534247-81813279 (*i.e*. ∼80 Mega base pairs), within the Omni 2.5 genotyping array framework. The data is formatted in the following way: rows are phased haplotypes; columns are positions of alleles represented by 0 when the corresponding nucleotide is identical with respect to the one in the reference genome (GRCh37) and 1 when it is point-wise mutated. Hence, the dataset is represented as a binary matrix.

### 2.3 PCA-WGAN

A Generative Adversarial Network (GAN) comprises two competing neural networks, the discriminator and the generator. The discriminator learns to dissociate real data from synthetic ones, while the generator aims at fooling the discriminator in a way that it fails at this classification task. Vanilla GANs face several problems, such as mode collapse, training instability, vanishing gradients, and sensitivity to hyperparameters (Arjovsky et al., 2017; Gulrajani et al., 2017). To account for these challenges, and following its successful application to population genetics (Booker et al., 2023; Yelmen et al., 2023), we used a Wasserstein GAN (Arjovsky et al., 2017) with gradient penalty (Gulrajani et al., 2017) (WGAN-GP) implemented in PyTorch (Paszke et al.). Whereas the discriminator of a vanilla GAN produces a probability of being fake or real, the critic of a WGAN produces an unbounded “realness” score for fake and real samples. The loss of a WGAN-GP, based on the estimation of the Earth Mover’s distance by the critic of two distributions on a batch of samples, is as follows :

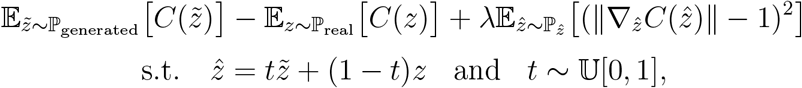

where *C* is the critic; 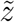 (resp. *z*) is a sample from the generator modeled distribution ℙ_generated_ (resp. from the data distribution ℙ_real_). The gradient penalty term, evaluated at a point 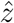 interpolated between a real and a fake point, is scaled by a constant factor *λ* and constrains the critic’s gradient to be close to 1. This gradient penalty regularization (Gulrajani et al., 2017) is a stabler alternative than weight clipping (Arjovsky et al., 2017), for enforcing the 1-Lipschitzianity property of the critic, a condition required by the Kantorovich-Rubinstein duality (Villani, 2009), otherwise called 1-Wassertein distance (𝒲_1_). 𝕌[0, 1] is the uniform distribution on [0, 1], yielding a distribution 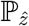 over 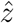. Enforcing the critic’s gradient to be of unit norm everywhere is intractable as it would require a number of samples exponential in the number of dimensions, instead (Gulrajani et al., 2017) showed that this property is met for an optimal critic along straight lines connecting pairs of points from both real and synthetic distribution.

The PCA-WGAN workflow combines an initial PCA on a binary matrix (5008 samples, 65K SNPs, see Section **2.2**), retaining 4507 components (90% PCs, see **Section 3.1** & **S4**) with a generative learning process in this PC space (**FIG.1.A**). To generate new fake binary SNP data, we sampled points in the space learned by the GAN (modeling the PC space), then performed the PCA inverse transformation using eigenvectors computed from real data PCA, and applied a binarization step based on a threshold of 0.5 (see Supplementary text **S4** and Section 2.6 for more details and alternative procedures).

### 2.4 Glocal-PCA-WGAN: Chaining multiple local PCA-WGAN

We designed an improvement of PCA-WGAN, called Glocal-PCA-WGAN (G stands for global), that explicitly leverages both global and local genomic scales. To circumvent difficulties arising from performing dimension reduction in the 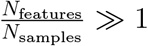 regime, our approach segments the SNP matrix into *K* multiple blocks. The procedure goes as follows: we split the SNP matrix into three successive blocks of equal number of SNPs and applied PCA to each block (i.e., each genomic region) independently of each other (yielding the colored blocks labeled 1, 2 and 3 in **FIG. 1B**). In this architecture, there are multiple local critics and one global critic. Each local critic aims to capture the local information of its assigned block, while the global critic should recover the relations between the blocks, i.e., at a larger genomic scale. The WGAN-GP is trained in the space of the concatenated PCA projections. Once the training is done, we sampled the generator, split the generated PC scores into three parts and inversed transform each chunk with the corresponding PCA (see Appendix **S5** for more details).

**Figure 1:**
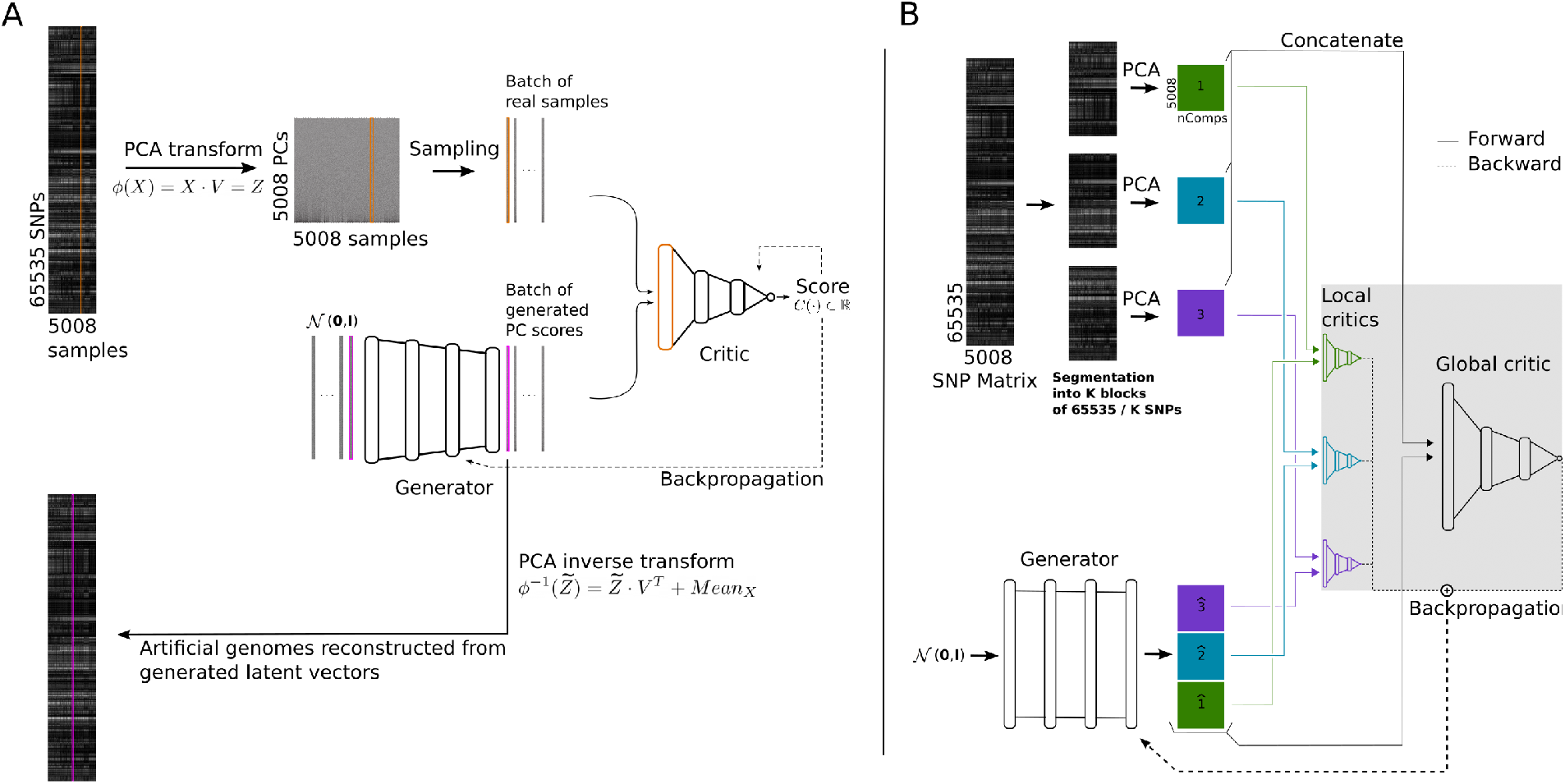
PCA-WGAN architectures. **A**. PCA-WGAN. The SNP matrix of 5008 samples by 65535 SNPs is reduced via PCA to a PC score matrix of 5008 samples by 4507 dimensions (90% PCs). The WGAN learns the PC space distribution. Once the training of the WGAN generator and critic is finished, new PC scores are generated and mapped back to the data space via PCA inverse transform. **B**. Glocal-PCA-WGAN. The SNP matrix is split into even-sized chunks, to which PCA is independently applied. Each local critic is fed with a single PCA chunk (colored block labeled 1, 2, or 3) and a generated PC scores chunk (colored block labeled 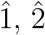 or 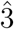), while the global critic is fed with the concatenated PCA projections and generated PC scores (i.e. all colored blocks).

#### Constraints on the choice of block number *K*

Increasing *K* leads to an increased difficulty for the WGAN as it will not only have to recover long-range interactions between the blocks but also short-range interactions since increasing *K* leads to higher risk of breaking interactions within an LD block. Furthermore, if *K* is high enough such that the blocks have fewer features than samples, then either (i) keeping all principal components amounts to no reduction at all (i.e., the output size of the generator & input size of the global critic will be 65K), or (ii) keeping fewer components leads to substantial reconstruction errors. On the other hand, if *K* is too small, i.e., the genomic chunks are large, we are compressing too many SNPs into *N*_samples_ dimensions. This is complicated further by the fact that keeping all PCs yields an input size of *K × N*_samples_ for the global discriminator, which should be kept low. All in all, we want *K* to be small but not so much so that PCA blocks still capture local information. Hence we chose *K* = 3 to keep the balance between reasonable parameter size and retaining a number of PCs that still yields a nearly optimal reconstruction error.

### 2.5 Light PCA-WGAN

Finally, we designed a light version of PCA-WGAN. This light model learns to mimic the principal components displaying multi-modal structure, i.e. the first six PCs, and is associated with a counterpart that models the remaining 5002 PCs thanks to a multivariate normal distribution. We trained a neural network (NN) to predict the parameters of the multivariate normal distribution (**FIG. S9**) and coined this model as Light PCA-WGAN with NN. Precisely, the neural network predicts the means and variances of these remaining PCs (low variance) from the first six PCs (high variance). The neural network is optimized on the real training data through the minimization of the Gaussian negative log likelihood loss (Nix and Weigend, 1994).

#### Sampling

To create novel genomic sequences, we first generated 6-dimensional samples with the Light PCA-WGAN. We then sampled the low variance PC scores conditionally on these high variance PC scores. For this, we ran the 6-dimensional samples through the NN to recover the parameters of a diagonal multivariate normal distribution and sampled this distribution. Finally, we concatenated the high and low variance parts of a sample and inverse-transformed it with PCA.

### 2.6 Binarization

Latent generative modeling requires projecting synthetic samples from the latent space to the data space once the samples are generated. Given that PCA is not designed for discrete spaces, it assumes continuity for the data space, resulting in inverse transformed samples living in a continuous space. Consequently, a final transformation is necessary to map these samples back into SNP space. We evaluated two approaches for this transformation: a naive binarization method applying a fixed threshold of 0.5 for all sites, and a tailored binarization approach with site-specific thresholds.

For that latter approach, at each site we computed the cumulative distribution of the reconstructed synthetic samples and the allelic frequency in the real data of that site. The threshold was then obtained by running the frequency of the real data through the inverse cumulative (derived by linear interpolation of the empirical cumulative distribution).

### 2.7 Shrinkage correction

We split the SNP matrix into a train and test sets of equal size, applied PCA to the training set, and used the resulting basis of eigenvectors to project test samples that were not included in the initial PCA. In the high dimension low sample size regime (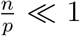 where *n* is the sample size and *p* is the number of features), a shrinkage phenomenon appears, as described by Lee et al. (2010, 2014). This phenomenon is related to the asymptotic behavior of the sample eigenvectors, indicating a discrepancy between the population and sample eigenvectors.

Following a correction proposed by Lee et al. (2010) for PCA projection bias, we adjusted each PC score of a shrunk distribution *D* (either the synthetic distribution from a generative model or the projected test set) with a multiplication factor equal to the inverse empirical shrinkage factor, 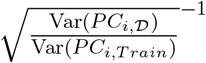, where Var(*PC*_*i,D*_), Var(*PC*_*i,T rain*_) are the variances on the i-th PC of the generated distribution in the latent space or the projected test set, and the projected train set respectively.

### 2.8 Sensitivity of 𝒜𝒜_*T S*_ with respect to shrinkage

𝒜 𝒜_*T S*_ (Yale et al., 2020) is a score based on nearest neighbors to assess overfitting for the synthetic data, which can further be used to derive a privacy score by comparing the resemblance of synthetic to train and test datasets.

This score compares the distances between pairs of samples within a distribution, as well as between distributions. Namely, *d*_TT_(*i*) is the distance of a fixed real sample indexed by *i* to its nearest neighbor in the real data, while *d*_TS_(*i*) is the distance of a fixed real sample indexed by *i* to its nearest neighbor in the synthetic data. The distances *d*_SS_(*i*) and *d*_ST_(*i*) are defined similarly. In an ideal scenario of perfect resemblance between the real and synthetic distributions, the nearest neighbor of a real sample would be equally likely to be synthetic as real, and similarly, the nearest neighbor of a synthetic sample would be equally likely to be real as synthetic.

#### Definition

We assume a training set 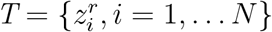 of size *N* (*r* for real), and a synthetic set of same size 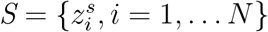

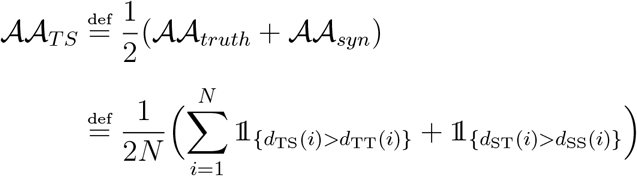

where

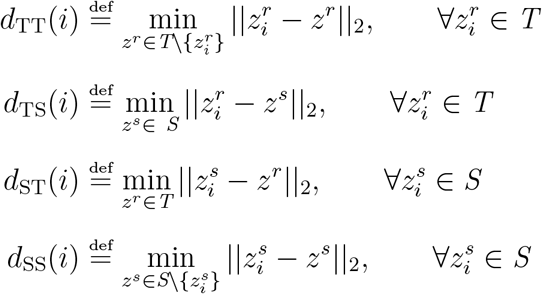

We further model the effect of shrinkage (see Subsection 2.7) on this indicator, *λ* being the shrinkage factor.

#### Proposition

Let *p*_*T*_ = 𝒩 (0, 𝕝_*D*_), *p*_*S*_ = 𝒩 (0, *λ 𝕝*_*D*_) with *λ* ∈]0, 1[ and *D* ∈ ℕ be the latent representation dimension.

Let 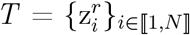 and 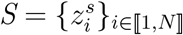, where 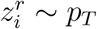 and 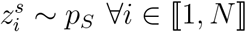.

Then

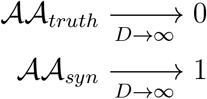

Our proposition illustrates that in a simple scenario of high-dimensional nested distributions, a generation anomaly often known as mode shrinkage (Stein et al., 2023; Yamaguchi and Fukuda, 2023; Montavon et al., 2016), 𝒜𝒜_truth_ converges to 0 and 𝒜𝒜_syn_ converges to 1, *i.e*. 𝒜𝒜_𝒯 𝒮_ converges to 0.5, yet the two distributions are easily distinguishable. Numerical experiments were conducted to confirm this assertion (see Appendix S3.6).

### 2.9 *k*-SNP motifs

We investigated the fine-grained structure of synthetic haplotypes. Specifically, we considered non-overlapping windows of size *k* along a haplotype, aiming to discern how generative models explore through these discrete spaces of sizes 2^*k*^. Inspired by *k*-mer commonly used in bioinformatics analyses, we dub *k*-SNP motifs such subsegments. For each motif of size *k*, with *k* ranging from 4 to 2048, we counted the number of unique *k*-SNP motifs within each non-overlapping window of the same size (**FIG.4.A**)

Secondly, we re-evaluated *k*-SNP motif counts after correcting for potential ancestry biases. As African ancestry contains higher genetic variations, a generative model that creates more samples of African ancestry than contained in the real dataset, might yield higher SNP motif diversity. We generated an excess number of AGs and then downsampled them to match the exact number of samples per continental ancestry group present in the real dataset. To achieve this, we labeled the AGs using a k-NN algorithm. Finally, we counted motifs within each continental ancestry group (**FIG.S12**).

A power-law was fitted on the distribution of *k*-SNP motif diversity for *k* ranging from 4 to 2048. To do that, we compiled the motif diversity for each *k* (the boxes in **FIG.4.A**.) into a single distribution, hence the information relative to *k* is abstracted away (**FIG.4.C**).

## 3 Results

As our latent generative modeling approach is based on learning in a reduced space, we first investigated in *Subsection 3.1* whether this smaller space retained enough information regarding the genomic space. In *Subsection 3.2* we describe shrinkage, a phenomenon that might appear both in PCA and in the generative case, albeit stemming from different causes.

Finally, we evaluated the artificial genomes generated with various models trained on a human dataset (The 1000 Genomes Project Consortium, 2010). Previously, Yelmen et al. (2021) investigated the quality of generated data, based on a set of summary statistics coming from population genetics. Here, we present a subset of these metrics and novel ones on the synthetic 65K SNPs data generated by PCA-WGAN, Glocal-PCA-WGAN, Light PCA-WGAN and compare these to previous WGAN and RBM models, which learn directly in the genetic space (Yelmen et al., 2023). The quality assessment of the synthetic data is presented in *Subsections 3.3, 3.4,3.5*.

### 3.1 Principal components space is a fertile ground for learning

We performed various pre-hoc training experiments to decide how many dimensions (i.e., PCs) should be kept after PCA and to gauge the errors caused by reconstruction. These experiments were performed considering a separate test dataset consisting of real genomes as the best-case generation outcome.

We first wanted to measure the quality of PCA encoding-decoding. Specifically, we aimed to determine whether the data generated in PC space can be reliably mapped back to the original space, assuming the generation step was successful. To check this, we randomly partitioned the dataset into a train and a test set. Both sets follow the same distribution; in that, the same modes appear in both, and they have similar densities. The test set acts as the perfect case where an ideal generative model completely captures the target distribution. Hence, a test set should give us hints on the expected behavior of properly generated data after reconstruction.

We considered the following protocol that does not rely on generated samples. First, we split randomly the dataset (*N*_samples_ = 5008) into a train and a test set of equal size and performed a PCA on the training set, varying the number of kept components. Then, we projected the test set onto the principal subspace derived from the training set (centered beforehand). Finally, we reconstructed the projected test set back into the original space and binarized it. Binarization was obtained through quantization with a threshold set to 0.5. We computed the mean reconstruction error of the binarized reconstructed test set with respect to the initial test set. The reconstruction error for one sample in the data space is defined as 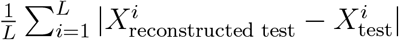 where *L* is the number of SNPs, 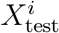 and 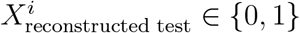. The reconstruction error for the training set was zero when keeping 100% of the PCs (**FIG.2.a**), *i.e*., there is no loss of information, but was equal to 0.06 in average for the test set. Decreasing the number of retained PCs reduced the gap between the train and test errors, indicating limited overfitting when few components were kept. However, because the test reconstruction error was at its minimum when keeping all components, we applied this final procedure for all remaining analyses.

To put this average test reconstruction error value (6%) into context, we computed the distribution of genetic distance separating two individuals (measured via Hamming distance, **FIG. 2.c**). Note that the reconstruction error applied to binary data is exactly the Hamming distance normalized by the sequence length, allowing direct comparisons. We found that the test reconstruction error (6%) is way below the minimal genetic distance in the considered dataset (13200, corresponding to ∼ 20% of errors along the SNP sequence), indicating that the error caused by reconstruction is smaller than the distance between the closest individuals.

**Figure 2:**
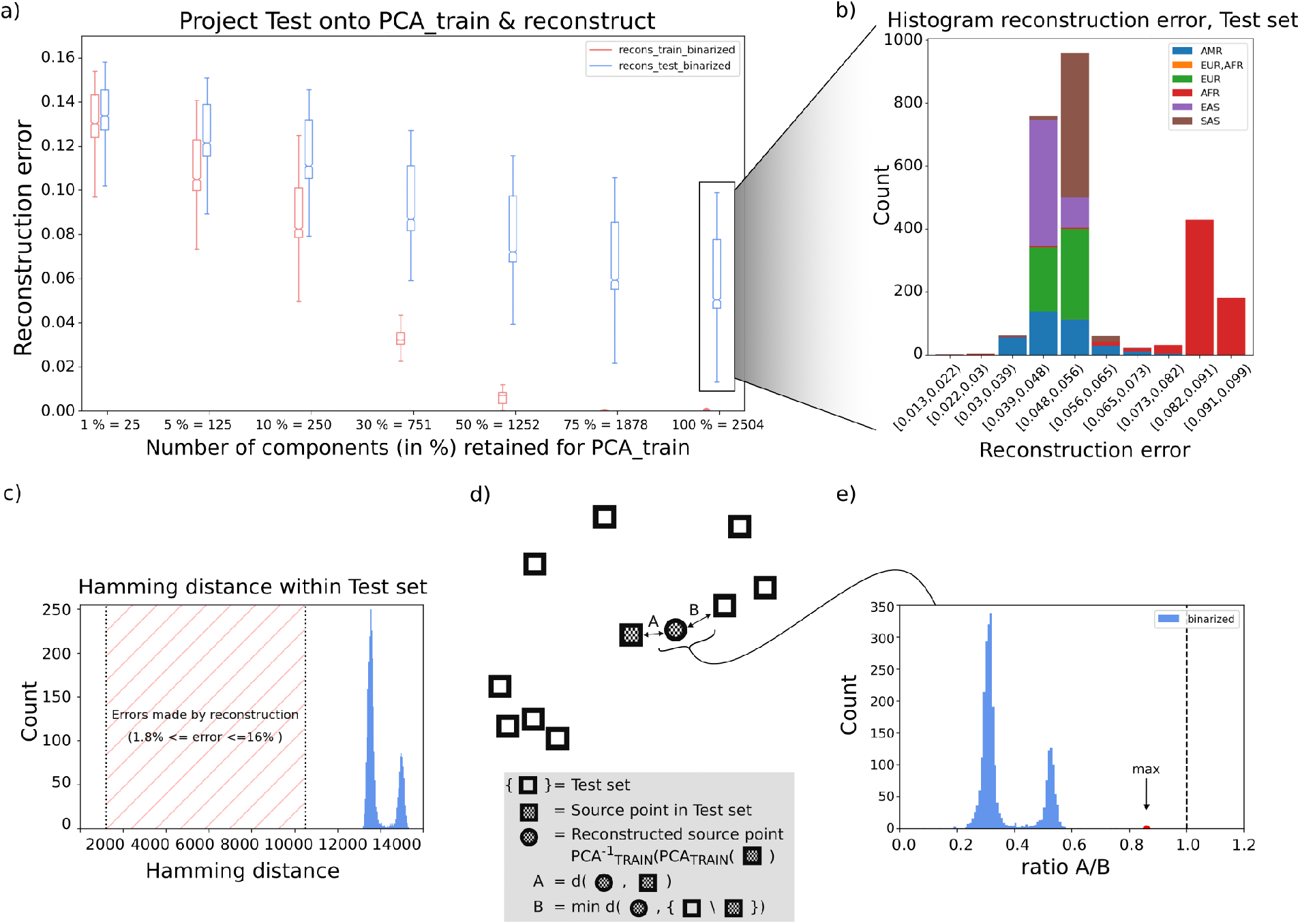
PCA reconstruction analyses. **a)** Reconstruction error as a function of the number of components retained when applying PCA to the training set. The reconstruction error is plotted after reconstructing the training set (red), and after projecting the test set onto the eigensubspace of the training set and reconstructing it (blue). **b)** Histogram of the reconstruction error for the reconstructed test set, keeping all components, colored by superpopulation labels. The superpopulation labels correspond to: AMR: American ancestry, EUR,AFR: European and African ancestry, EUR: European ancestry, AFR: African ancestry, EAS: East Asian ancestry, SAS: South Asian ancestry. **c)** Hamming distance within the test set. **d)** Illustration of how much reconstruction moves data points. **e)** Distribution of the ratio of the distance *A* between a source point and its reconstruction, by the distance *B* between the reconstructed source point and its nearest neighbor in the test set minus the source point.

We refined this analysis at the individual level by investigating, for each reconstructed point, whether its nearest neighbor is the source point (desired behavior) or another individual of the test set **(FIG. 2.d-e)**. Specifically, we compared the (Hamming) distance *A* (between a source point in the test set and its reconstructed point) to the distance *B* (between the reconstructed source point and its nearest neighbor in the test set excluding the source point). If *B < A* then the reconstruction step locally distorts the data to the extent that the information at the sample level is lost in the process, *i.e*., a reconstructed sample can be confused with another test sample through minor alterations. Such an unfaithful reconstruction would discourage to learn the generative model in the PC space.

We found that this was not the case **(FIG. 2.e)**, as the distribution of the ratio 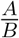 was considerably lower than 1 (dotted vertical line), with a max value equal to 0.82. This further demonstrated that the information content of the reconstructed source point remains consistent with that of the source point.

Interestingly, the distribution of the reconstruction error appears to be bimodal. We discovered that samples with a higher reconstruction error were of African ancestry while samples from the rest of the world were mixed in the first mode (**FIG. 2.b**). This is likely due to the known higher genetic diversity present on this continent (explainable by human past demographic history) (Tishkoff and Williams, 2002; Campbell and Tishkoff, 2008; The 1000 Genomes Project Consortium et al., 2015).

We further investigated whether the low variance principal axes were relevant when studying the test set (i.e., whether they are better than random directions). For this, we selected *n*_Comp_ principal axes and constructed the remaining 2504 − *n*_Comp_ dimensions as random directions, by sampling Gaussian vectors 𝒩_*k*_(**0, I**) with *k* = 65535 and orthonormalizing them with respect to the first *n*_Comp_ axes (Boyd and Vandenberghe, 2018). From **FIG. S2**, we observe that adding random directions to the *n*_Comp_ first principal axes decreased the reconstruction error as expected, since there is more information when considering more directions, i.e., when the projection space gets larger. However, adding the next principal axes instead of random directions decreased the reconstruction error substantially more. This confirmed that the low variance principal axes contained relevant information for the reconstruction step.

### 3.2 Shrinkage phenomenon in high dimensional low sample size setting

We performed an optimization-free experiment, similarly to Section 3.1, where the test set, that serves as synthetic set, is used to understand the expected behavior of a distance-based metric, 𝒜𝒜_*TS*_ (see Method 2.8), on generated samples.

After projecting the test set onto the basis obtained from PCA on a train set, we observed lower variances for PC scores of test samples than train samples (**FIG. S3**). This shrinkage phenomenon, as previously studied in (Lee et al., 2010, 2014), occurred in scenarios with high-dimensional data and low sample sizes. Specifically, it affected the PC scores of test samples not originally included in the PCA performed on a training set.

This phenomenon might lead to significant alterations in distances between the two sets in the PC space. As the density of the projected test set is shrunk with respect to PC scores of the train set (**FIG. S3**), the nearest neighbor of a projected test sample will mostly be a projected test sample rather than a train sample. Similarly, the nearest neighbor of a train sample will mostly be a projected test sample. That is, 𝒜𝒜_*syn*_ is close to one while 𝒜𝒜_*truth*_ is close to zero (see scores for projected and reconstructed test in **FIG. S5**).

To investigate the double impact of shrinkage and space dimension, we computed *AA* scores on different slices of the PC space, where shrinkage always exists but can be more or less prominent (**FIG. S3**.) In all cases, we observed that 𝒜𝒜*truth* and 𝒜𝒜*syn* converged towards 0 and 1, respectively, quite rapidly, *i.e*., generally for dimension 40 and higher, with a speed that depends on the expected shrinkage strength (**FIG. S4**).

To further confirm this effect of shrinkage on 𝒜𝒜_*T S*_, we applied a dilating factor as a correction (see Method 2.7). This correction led to a better behaved 𝒜𝒜_*T S*_ in PC space, where the different components were closer to the expected 0.5 value, as shown for projected test corrected on **FIG. S5**. It is worth mentioning here that 𝒜𝒜_*T S*_ in PC space or SNP space without binarization is the same since PCA is just an orthogonal transformation. However, we found that the binarization step with a fixed threshold set to 0.5 was deteriorating the 𝒜𝒜scores (see scores for reconstructed test corrected in **FIG. S5**).

Finally, we reproduced the conditions of the proposition (see Methods 2.8). We constructed two generic multivariate normal distributions 𝒩 (0, 𝕝_*D*_) (serving as real data) and 𝒩 (0, *λ* 𝕝_*D*_) (serving as synthetic data) with shrinkage factor *λ* ∈]0, 1[, *D* = 200 and generated 5000 samples from each distributions. We observed convergence of 𝒜𝒜_*truth*_ to 0 and 𝒜𝒜_*syn*_ to 1, where the speed of convergence is controlled by *λ*. For extreme shrinkage, *λ* = 0, there is a first regime where both 𝒜𝒜scores are equal to 1 (dark curves), then 𝒜𝒜_*truth*_ suddenly drops as the number of dimensions increases (see **FIG. S8**).

With these experiments in mind, we will systematically analyze the two 𝒜𝒜_*T S*_ components separately, as it allows to detect shrinkage on top of other generation anomalies (Yelmen et al., 2023).

### 3.3 Global patterns of genetic variation: principal component analysis and allelic frequencies

Real data exhibited a strong population structure pattern captured by PC1 to PC4 (**FIG. 3.a**), which was effectively replicated in AGs from all models, as can be seen through the 1D Wasserstein distance between the real and AG projections. Overall the error curve for PCA-WGAN was always below that of the RBM except for PC5, where the model was generating highly concentrated scores, yielding an increased Wasserstein distance. This concentration issue was not encountered in Light PCA-WGAN.

**Figure 3:**
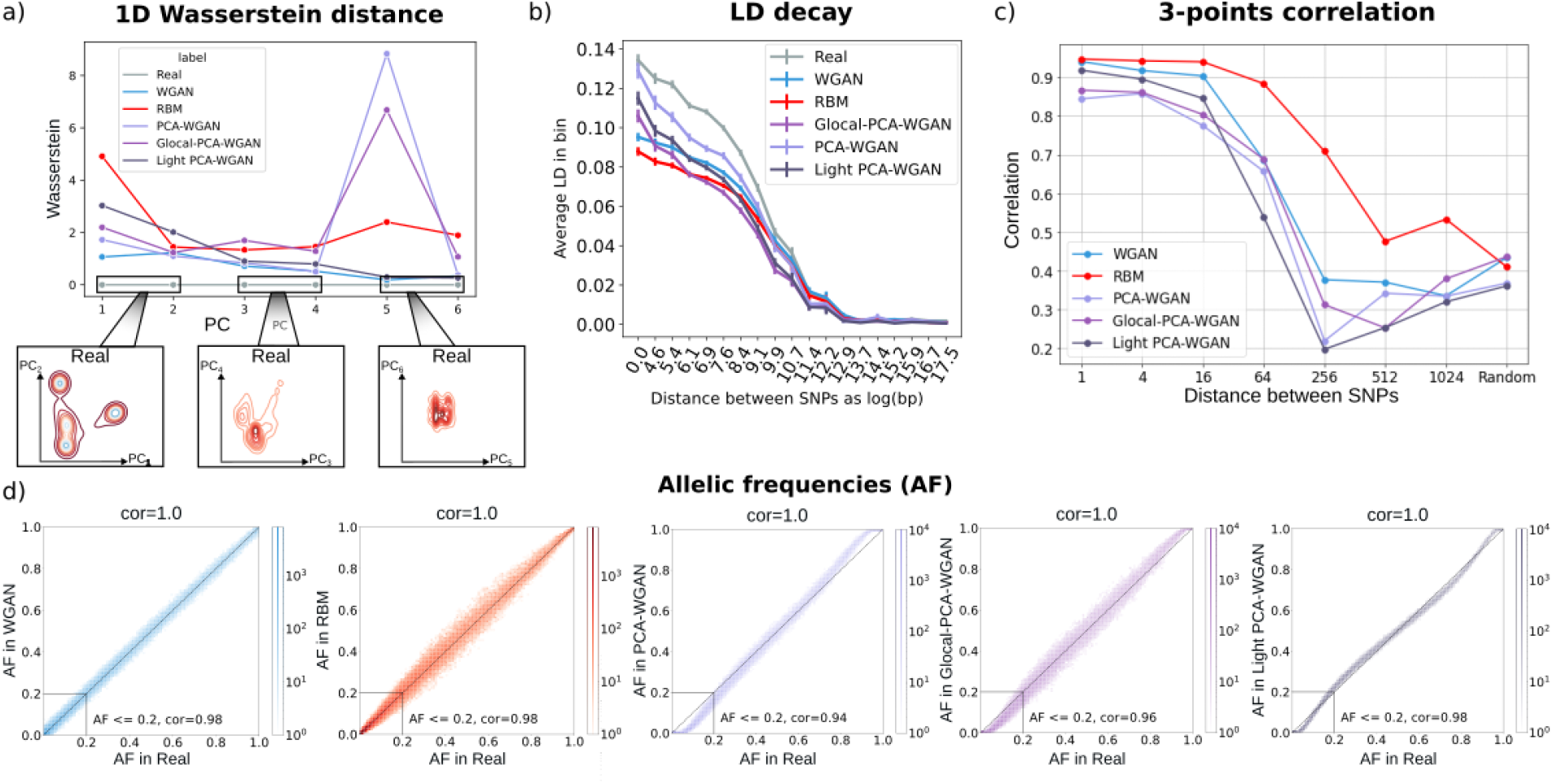
Population genetics summary statistics. **a)** The real and the synthetic data were concatenated into a single dataset, to which PCA is applied. The 1D Wasserstein distance is computed on each PCs. Real data is the grey curve, WGAN from (Yelmen et al., 2023) in blue, RBM from (Yelmen et al., 2023) in red, PCA-WGAN in light purple, Glocal-PCA-WGAN in dark purple and Light PCA-WGAN with NN in grey. The density plot of real data for the first six PCs is detailed in first column of **FIG.S13. b)** LD decay approximation. **c)** Three-points correlation separated by a varying amount of SNPs. **d)** 2D Histogram of the allelic frequencies in real data (x-axis) and in AGs produced by the models.

The synthetic datasets followed overall the same trends as the real one, in terms of allelic frequencies (**FIG. 3.d**). For frequencies ≤ 0.2, the correlation score between real and AGs ranged from 0.94 to 0.98. Nonetheless, PCA-WGAN fixed many alleles (20523 ∼ 31% of the sites; 15016 for Light PCA-WGAN; 12686 for Glocal-PCA-WGAN; 2672 for WGAN; 55 for RBM; 8 in real), especially the ones which are rare in the original dataset, similar to previous findings from different models (Yelmen et al., 2021, 2023).

### 3.4 Local haplotypic structure: zooming on multi-locus interactions

The linkage disequilibrium (LD) curve shows how the correlation between pairs of SNPs decreases as a function of their physical distance (**FIG. 3.b**). PCA-WGAN reproduced these statistics better than all the other models, *i.e*., the LD curve was the closest to the real curve, notably for short-range interactions. The lower LD observed for GlocalPCA-WGAN was likely due to the uniform splitting of the SNP matrix (yielding blocks with the same number of SNPs). A potential future improvement in this regard might be to partition into regions that preserve LD blocks (Privé, 2021) (e.g., by splitting at recombination hotspots). Finally, Light PCA-WGAN fell in-between PCA-WGAN and Glocal-PCA-WGAN, and still yielded more accurate short range interactions than WGAN and RBM, even though it has one to two orders of magnitudes fewer parameters (**Table 1**).

**Table 1:**
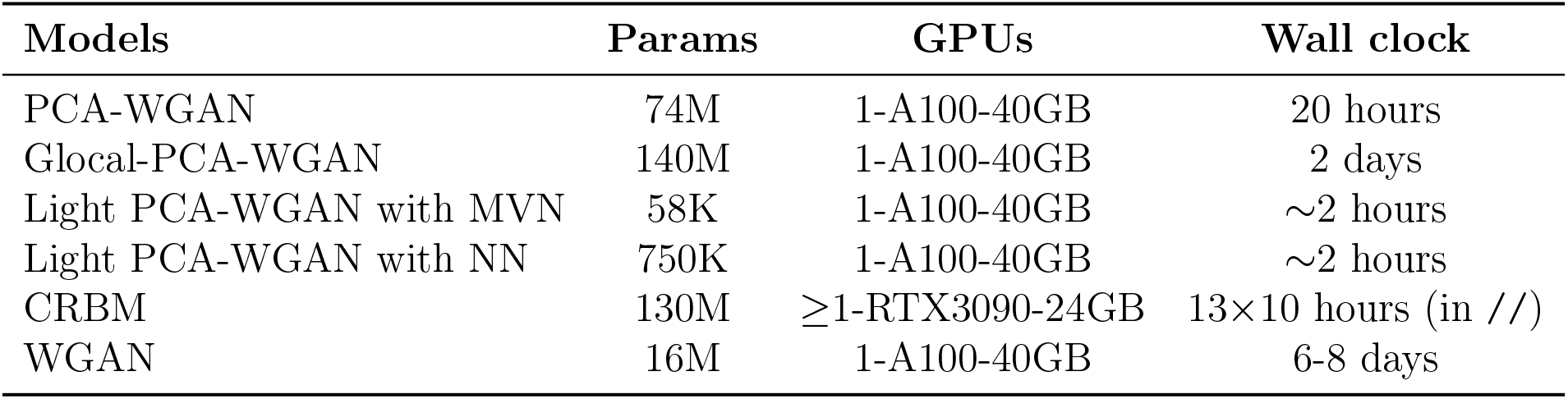
Training GPU & runtime comparison between models for 65K SNP dataset.

| Models | Params | GPUs | Wall clock |
| --- | --- | --- | --- |
| PCA-WGAN | 74M | 1-A100-40GB | 20 hours |
| Glocal-PCA-WGAN | 140M | 1-A100-40GB | 2 days |
| Light PCA-WGAN with MVN | 58K | 1-A100-40GB | $\sim 2$ hours |
| Light PCA-WGAN with NN | 750K | 1-A100-40GB | $\sim 2$ hours |
| CRBM | 130M | $\geq 1$-RTX3090-24GB | $13\times 10$ hours (in *//*) |
| WGAN | 16M | 1-A100-40GB | 6-8 days |

In addition to LD, we computed the three-point correlation statistics for SNP triplets separated by *1, 4, 16, 64, 256, 512, 1024* and a random number of SNPs respectively (**FIG. 3.c**). While PCA-WGAN had lower correlations than RBM model in Yelmen et al. (2023), it remained close to the convolutional WGAN model, and demonstrated similar results to both models in the challenging scenario involving SNP triplets separated by random distances. Multiple local critics (Glocal-PCA-WGAN) improved this statistic for almost all SNP distances (except for a distance of 512 SNP) and even marginally surpassed WGAN and RBM benchmark models for SNPs separated by random distances. Light PCA-WGAN performed better than PCA-WGAN and Glocal variant for short range interactions but its performance declined as the distance between SNPs increases.

After evaluating data quality with summary statistics proposed previously (Yelmen et al., 2021, 2023), we introduced a new one, based on SNP motifs. This allowed us to dissect more closely whether local haplotypic patterns are well preserved, which was previously mostly deduced from the 3-point correlation results. Artificial genomes unanimously exhibited higher motif diversity than in real data (see Method 2.9 and **FIG.4.A**.). For *k*-SNP motifs with *k* ≤ 16, AGs from PCA-WGAN had a diversity lower than other generative models and closer to the real data, which is in line with LD of PCA-WGAN being the closest to real LD for smaller SNP distances. Indeed, highly correlated SNPs tightly constrain how the motif space should be populated. The diversity pattern shown in **FIG.4.A**. was maintained when we focused on each population independently, i.e., AGs had higher diversity than real sequences and those from PCA-WGAN were closer to the real data for *k* ≤ 16 (**FIG.S12**).

We further fitted a power-law to the distribution of *k*-SNP motif diversity for *k* ranging from 4 to 2048, and retrieved the exponent as a measure of complexity of the data. A lower power-law exponent indicates higher diversity, i.e. more unique motifs. PCA-WGAN had the lowest exponent, which corresponds to a longer tail, while WGAN and RBM displayed a multimodal distribution, where bumps corresponds to particular *k*-SNP motif sizes (**FIG.4.C**.), i.e., these two latter models produced a diversity that was homogeneous along the haplotype windows. While *k*-SNP motif hallucination seemed to be a salient feature of generative models (**FIG.4.A**.), PCA-WGAN and its alternative in fact possessed (through the power law exponent) the highest diversity (**FIG.4.C**.). *k*-SNP motifs of size *k* ≥ 32 were likely contributing to this inflated diversity for PCA-WGAN and its Light variant compared to other models.

We further computed Shannon’s entropy over each window of size *k* and compared it to a Bernoulli model having parameters equal to the frequency at each site of the real data. Note that for random *k*-SNP motifs drawn from a uniform distribution, the entropy will be equal to *k*. Since a *k*-SNP motif can take on 2^*k*^ possible states, and our real and synthetic datasets provided 5008 samples per window, the distribution might not be well sampled for *k* ≥ 12, *i.e*., when there are more than 2^12^ = 4096 states. However, rare *k*-SNP motifs that are missed would not have significantly affected the entropy, as long as they are not too numerous. Entropy calculations showed that a substantial amount of local haplotypic structure was present in real and AGs data compared to random sequences (from the Bernoulli model). Furthermore, *k*-SNP motifs in AGs were slightly less structured than in real genomes (**FIG. 4.B**.).

**Figure 4:**
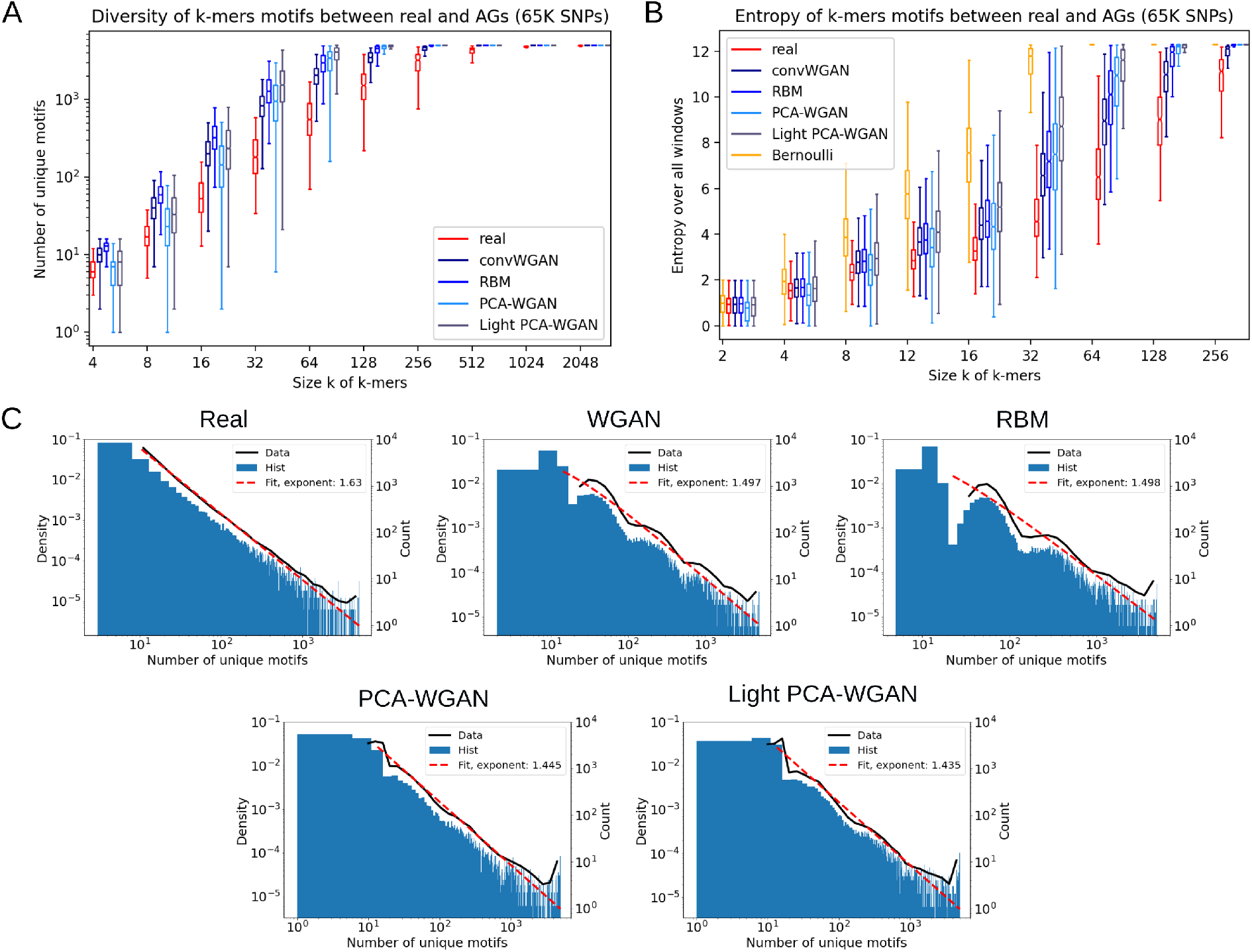
Diversity & entropy of *k*-SNP motifs. **A**. Distribution of the number of unique motifs over all non-overlapping windows of size *k*. Diversity of *k*-SNP motif per window in real data (red), and the synthetic counterpart (blue) for WGAN, RBM (Yelmen et al., 2023), PCA-WGAN and Light PCA-WGAN. **B**. Shannon’s entropy computed over all non-overlapping windows of size *k*. The Bernoulli model has parameters equal to the frequency at each site of the real data. **C**. Histogram of the diversity of *k*-SNP motifs for multiple *k*, with the probability density function as black curve and power-law fit as dashed red line. The power-law exponent is 1.63, 1.497, 1.498, 1.445, 1.435 for Real, WGAN, RBM, PCA-WGAN and Light PCA-WGAN respectively.

### 3.5 Measuring overfitting in generative models

We evaluated how well the AGs coming from the different models perform in terms of resembling real data (**FIG. 5**). Coincidentally, the 𝒜𝒜_*T S*_ score of PCA-WGAN and glocal variant showed a behavior similar to the one described for the test set, *i.e*., 𝒜𝒜_*syn*_ close to one and 𝒜𝒜_*truth*_ close to zero (see Subsection 3.2). This is likely due to the exceedingly narrow density of the generated points along the late PCs. However, unlike the shrinkage observed in the projected and reconstructed test set, this effect is most likely driven by the Wasserstein distance favoring shrunk modeled distributions (see Section 4.3 and FIG.5 in Montavon et al. (2016)). After correcting for the undesired shrinkage effect (see Method 2.7) the score improved not only over the PCA-WGAN but also over the WGAN and RBM from (Yelmen et al., 2023). This enhancement of 𝒜𝒜_*T S*_ came at the cost of slightly worse LD statistics, staying nonetheless above that of WGAN and RBM for short range interactions (**FIG. S7 Top right**). On the other hand, it also alleviated the high Wasserstein distance to the real AGs on PC5 and the allele fixing issue (**FIG. S7 Top left and Bottom**). Finally, Light PCA-WGAN palliated the difficulties arising from training with Wasserstein loss in high-dimensional space, with 100*×* fewer parameters than PCA-WGAN (**Table 1**), and achieved the best 𝒜𝒜_*T S*_ score among all models.

**Figure 5:**
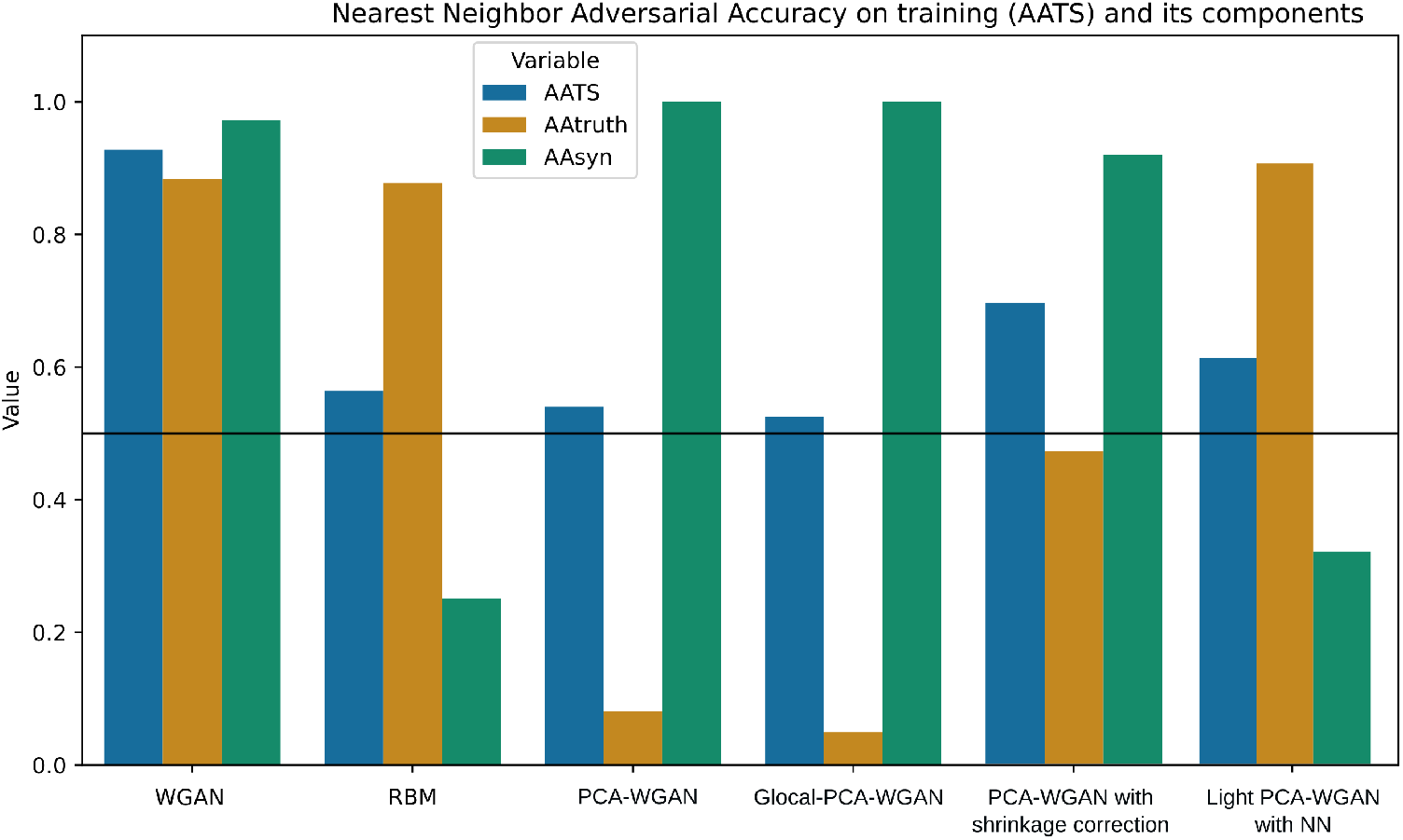
Nearest neighbor adversarial accuracy. 𝒜𝒜_*T S*_ score for AGs was computed in the data space. The black line indicates the optimum value whereas values below the line indicate overfitting and values above the line indicate underfitting.

Dissecting the 𝒜𝒜_*T S*_ of PCA-WGAN generated samples in PCA space, *i.e*. before their transformation into the genomic space, we also found imbalanced 𝒜𝒜_*syn*_ (close to 1) and 𝒜𝒜_*truth*_ (close to 0) scores. This further confirmed that the shrinkage effect was due to Wasserstein training as previously explained. However, 𝒜𝒜_*T S*_ of Light PCA-WGAN with NN in PCA space displayed preferable behavior w.r.t resemblance (**FIG. S6**). Similarly to Subsection 3.2, we identified that the binarization step with a fixed threshold set to 0.5 is deteriorating the 𝒜𝒜_*T S*_ score (**FIG.S6**). In an effort to circumvent both issues of allelic fixation (demonstrated in Subsection 3.3) and poor 𝒜𝒜_*T S*_, we applied a frequency-based binarization (see Methods 2.6) instead of a hard threshold. This frequency-based binarization yielded allelic frequencies that were identical w.r.t real data, yet the 𝒜𝒜_*T S*_ behaved badly, i.e., 𝒜𝒜_*T S*_ ∼ 0.5, 𝒜𝒜_*truth*_ ∼ 1 and 𝒜𝒜_*syn*_ ∼ 0. Hence, a fixed threshold set to 0.5 was preferred.

### 3.6 Computational efficiency comparison

We compared training compute resources between different models (**Table 1**), as well as the number of parameters as a function of sequence length (**FIG.S11**). The CRBM in (Yelmen et al., 2023) consisted of 13 RBMs with 10M parameters each, where each RBM was trained during ∼ 10 hours (wall clock time). The training time of the whole set of RBMs depends on the number of available GPUs, and can benefit from parallelization (//), *i.e*., with a sufficient number of GPUs, the training time can be at best ∼ 10 hours. While PCA-WGAN had more parameters than WGAN, its training time remained almost an order of magnitude smaller. Glocal PCA-WGAN had the highest number of parameters, but still was faster to train than WGAN. Light PCA-WGAN with NN had 100*×* fewer parameters than PCA-WGAN and 20*×* fewer parameters than WGAN, with the fastest training time, on par with Light PCA-WGAN with a multivariate normal distribution (MVN). That last model (see *Appendix S7*) provided similar results on the population genetics summary statistics and 𝒜𝒜_*T S*_ metric as Light PCA-WGAN with NN, only requiring 58K parameters, *i.e*., 1275*×* less than PCA-WGAN or 275*×* less than WGAN.

## 4 Discussion and conclusion

This study tackled scaling generative models for high-dimensional SNP data with PCAWGAN and its derivative models, and assessed AG quality through population genetics summary statistics, privacy-related scores and information theoretic metrics. To our knowledge, while dimension reduction is routinely used as a ML preprocessing step, its application in generative modeling for genomics is unexplored, making our reduced-space genomic data generation simple and innovative. Moreover, no other studies benchmarked their models on a very high-dimensional tabular dataset, except for models in (Yelmen et al., 2023) and the models presented in this current work. For example, Dang et al. (2023) demonstrated good performances for the 805 SNPs dataset (Colonna et al., 2014; Yelmen et al., 2021) but weaker ones for 10K SNPs, while *DNAGPT* Zhang et al. (2024) demonstrated performances comparable to (Yelmen et al., 2023) for 10K SNPs but did not scale up to 65K SNPs. Despite PCA-WGAN having a drawback in allelic frequencies, it achieved results comparable to recent advances (Yelmen et al., 2021, 2023) and even captured two-point SNP correlations (LD) better. On the other hand, Glocal-PCA-WGAN improved the allele fixing issue since the PCA compression is lighter, but produced worse results in terms of LD. This could potentially be improved by finding optimal splits for blocks with respect to LD (Privé, 2021). Furthermore, Glocal-PCA-WGAN could scale to longer SNP sequences. As we showed that PCA-WGAN on 65K SNPs offered competitive results, we could envision a Glocal-PCA-WGAN processing 3*×*65K SNPs. In this case, keeping 40% of PCs for each block would lead to the same parameter size as in the current Glocal-PCA-WGAN for 65K SNPs (**FIG.S5**). Finally, we showcased Light PCA-WGAN as a viable frugal alternative to complex sequence modeling architectures, obtaining similar results to large models with order of magnitudes fewer parameters.

As generative modeling becomes more widespread in various domains from natural language processing to image synthesis, defining the desired properties of synthetic data becomes more important (Alaa et al., 2022; van Breugel et al., 2023; van Breugel and van der Schaar, 2023). However, generative models often suffer from multiple shortcomings, one of them, *mode shrinkage*, consists in the concentration of the synthetic data on the modes of the real data without covering the full range (Stein et al., 2023; Yamaguchi and Fukuda, 2023; Montavon et al., 2016). Despite the effectiveness of shrinkage correction and Light PCA-WGAN in addressing the nearest neighbor adversarial accuracy issue, these experiments underscored the significance of distinguishing 𝒜𝒜_*truth*_ from 𝒜𝒜_*syn*_ components within 𝒜𝒜_*T S*_. This decomposition becomes particularly crucial as it reveals instances where the overall adversarial accuracy (𝒜𝒜_*T S*_) appears satisfactory, while individual components 𝒜𝒜_*truth*_ and 𝒜𝒜_*syn*_ exhibit poor performance. In addition, we showed that 𝒜𝒜_*T S*_, being distance-based, may lead to spurious results under certain circumstances, in particular when there are generation anomalies (Yelmen et al., 2023), over- or under-dispersion of samples due to shrinkage (*Proposition 2.8*), mode collapse, or drops in diversity. The adjustment of the PC scores via a dilating factor improved 𝒜𝒜_*T S*_ at the cost of slightly lowering the quality of AGs is sub-optimal and could be refined (see Method 2.7). Indeed, the non alignment of eigenvectors across train and test datasets induces a dilation in inaccurate directions. Finally, we observed that different set of hyperparameter values for the NN paired with Light PCA-WGAN (Figure S9) yielded substantially different results for *AA*_*T S*_. Instead of manual or grid search for the best set of hyperparameter values (described in Supplementary material S6), AutoML algorithms could be employed in future research.

Analysing the *k*-SNP motifs along a haplotype revealed simple yet discriminative feature between real and synthetic data. Namely, given a fixed window of size *k*, AGs contained significantly more motifs than in real data. Despite having their discriminative features, AGs were found useful for multiple tasks (e.g. imputation, selection scan, or ancestry inference) (Yelmen et al., 2021; Montserrat et al., 2019). This raises the question of balancing overall data quality with its utility for specific applications. Currently, there is no universally accepted indicator to effectively evaluate this balance, underscoring the need for further methodological developments. Indeed, classification algorithms that disentangle empirical from simulated data have recently highlighted the unrealistic nature of sequences generated by evolutionary models (Trost et al., 2023). However, there remains a lack of methodology to gauge the practical utility of these recognizable synthetic data.

Overall, PCA-WGAN is a promising novel methodology that combines simple dimensionality reduction and generative modeling to produce diverse and longer artificial genomes, making it a potentially valuable tool for enhancing genome-wide analyses with synthetic data, especially when access to real datasets is limited.

## 5 Declarations

### 5.1 Ethics approval and consent to participate

Not applicable

### 5.2 Consent for publication

Not applicable

### 5.3 Availability of data and materials

All the relevant data and code is publicly available at https://gitlab.inria.fr/ml_genetics/public/artificial_genomes/-/tree/master/1000G_real_genomes for data and https://gitlab.inria.fr/ml_genetics/public/artificial_genomes/-/tree/master/PCA-WGAN for the models.

### 5.6 Authors’ contributions

A.S : Conceptualization, Formal analysis, Investigation, Methodology, Project administration, Resources, Software, Validation, Visualization, Writing – original draft.

C.F : Methodology, Writing – review & editing.

G.C: Conceptualization, Methodology, Project administration, Supervision, Resources, Writing – review & editing.

B.Y : Conceptualization, Data curation, Formal analysis, Investigation, Methodology, Project administration, Supervision, Software, Writing – review & editing.

F.J : Conceptualization, Data curation, Funding acquisition, Methodology, Project administration, Resources, Software.

## 5.7 Acknowledgements

This work benefited from Inria TAU computing resources. We thank Michèle Sebag for insightful discussions.

