## Supplementary material for "Latent generative modeling of long genetic sequences with GANs"

#### S1 Overview

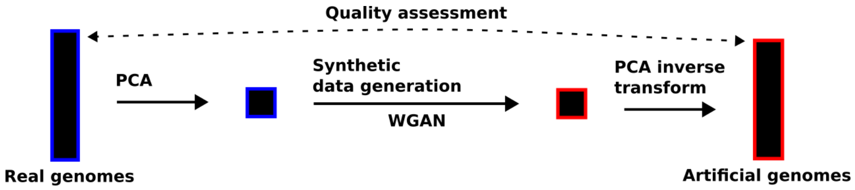


Figure S1: **Overview of the general method.**

#### S2 Reconstruction with PCs augmented by random axes

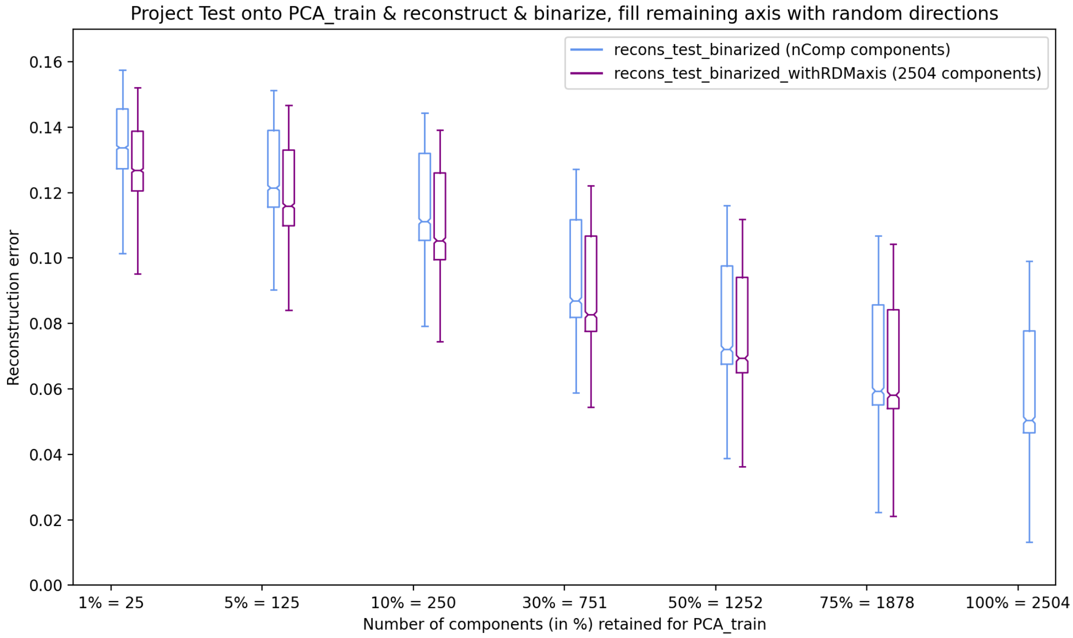


Figure S2: **PCA reconstruction analyses with random orthonormal axes.** The reconstruction error was computed for two procedures: (1) PCA was applied on the training data while varying the number of kept axes (x-axis), the test data was then projected onto the resulting eigensubspace, reconstructed and binarized (blue); (2) PCA was applied on the training data while varying the number of kept axes (x-axis), the remaining axes are filled with random gaussian vectors that are orthonormalized to the kept PCA eigenvectors and to themselves via Gram-Schmidt algorithm, the test data was then projected onto the resulting eigensubspace, reconstructed and binarized (indigo). Adding random directions decreased the reconstruction error, but not as much as adding remaining PCA modes.

.

### S3 Privacy metric biased by shrinkage

#### S3.1 Shrinkage

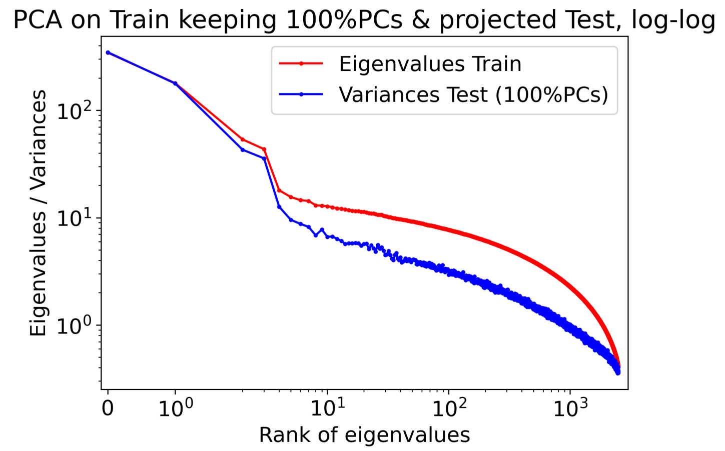


Figure S3: **Variance of the PC scores.** PCA was applied on a training set (2504 samples, 65535 SNPs) with all components kept. The test set (2504 samples) was projected onto the resulting eigensubspace. The variance of the PC scores of the train set (red) was higher than that of the test set (blue) due to a shrinkage phenomenon (see Subsection 3.2).

#### S3.2 $\mathcal{AA}_{TS}$ in PC space of a train-test split

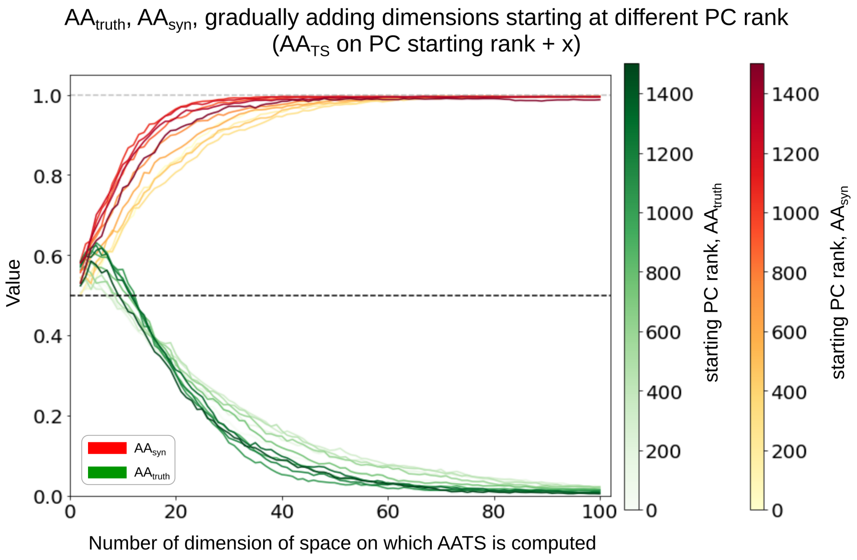


Figure S4: **$\mathcal{AA}_{TS}$ of the projected test set.** PCA was applied on a training set (2504 samples, 65535 SNPs) with all components kept. The test set (2504 samples) was projected onto the resulting eigensubspace. Starting from a chosen PC rank $s$, we gradually added dimensions along the $x$-axis and computed $\mathcal{AA}_{truth}$ (green), $\mathcal{AA}_{syn}$ (red) for each space, continuing until reaching 100 dimensions. That is, $\mathcal{AA}_{TS}$ is computed on $\text{Proj}_{\text{TEST}}[:, s : s + x]$, where $\text{Proj}_{\text{TEST}}$ represented the projected test set. The darker the curve's color, the higher the rank $s$ of the selected PC. The shrinkage strength is expected to be higher for low variance PCs as can be seen by the gap between red and blue curve in **FIG. S3**.

#### S3.3 Measuring the effect of shrinkage correction on $\mathcal{AA}_{TS}$

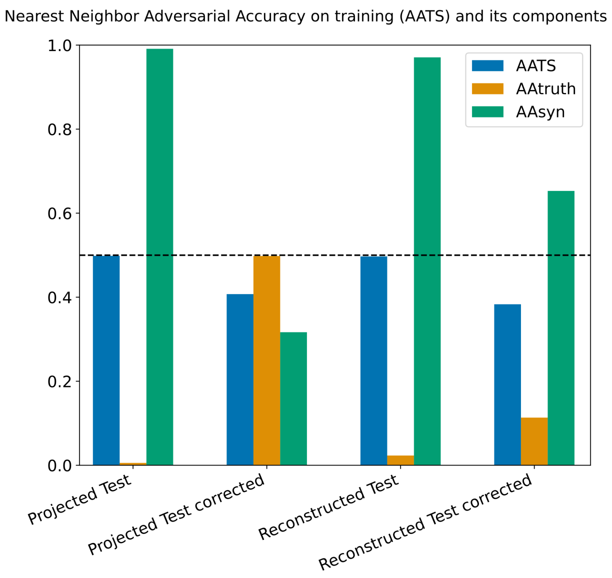


Figure S5: $\mathcal{AA}_{TS}$ **of the test set with shrinkage correction.** PCA was applied on a training set (2504 samples, 65535 SNPs) and all components are kept (i.e. 2504). The test set (2504 samples) was projected onto the resulting eigensubspace. A copy of the projected test set was made, and every PC scores were multiplied by the inverse of the empirical shrinkage factor (see Subsection 2.7). $\mathcal{AA}_{TS}$ was computed on the PC scores of the projected test set, the corrected version and their reconstruction, with the euclidean and hamming distance respectively.

#### S3.4 Aftermath of binarization on $\mathcal{AA}_{TS}$

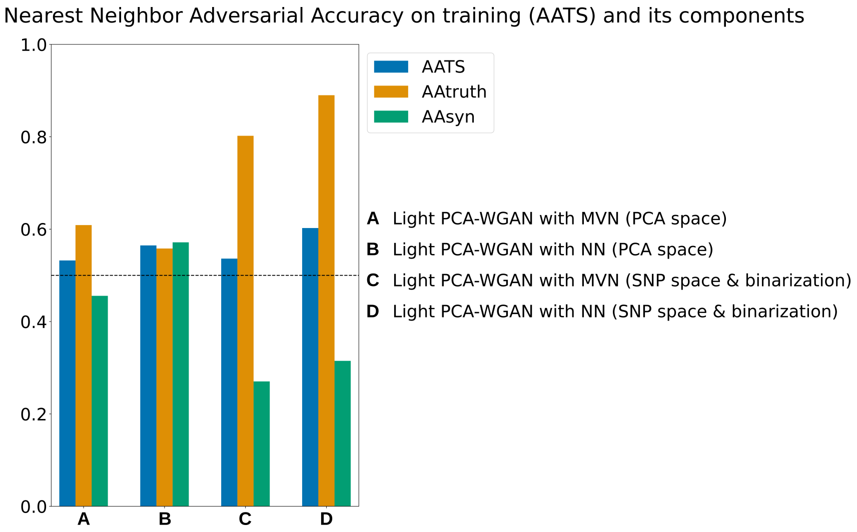


Figure S6: **Effect of binarization on** $\mathcal{AA}_{TS}$**.** $\mathcal{AA}_{TS}$ (blue), $\mathcal{AA}_{truth}$ (yellow), $\mathcal{AA}_{syn}$ (green) for AGs from Light PCA-WGAN with MVN and NN, computed in PCA space and SNP space with binarization step.

#### S3.5 Shrinkage corrected AGs

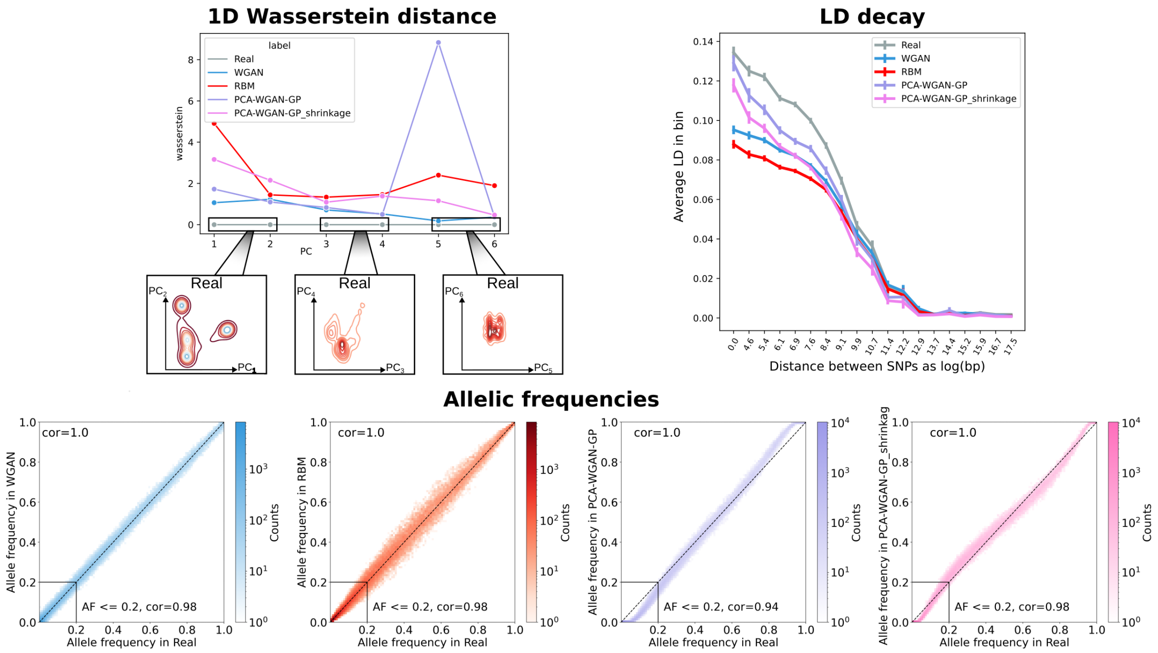


Figure S7: **Population genetics summary statistics. Top left:** The real and synthetic data were concatenated into a single dataset on which PCA was applied. The 1D Wasserstein distance between real and synthetic data was computed for each PC. Real data is displayed in grey, WGAN from (Yelmen et al., 2023) in blue, RBM from (Yelmen et al., 2023) in red, PCA-WGAN in light purple, shrinkage corrected PCA-WGAN in violet. **Top right:** LD decay approximation (correlation for pairs of SNPs as a function of their physical distances). **Bottom:** 2D Histogram of the allelic frequencies in real (x-axis) and in AGs produced by the models.

#### S3.6 Numerical validation of proposition 2.8

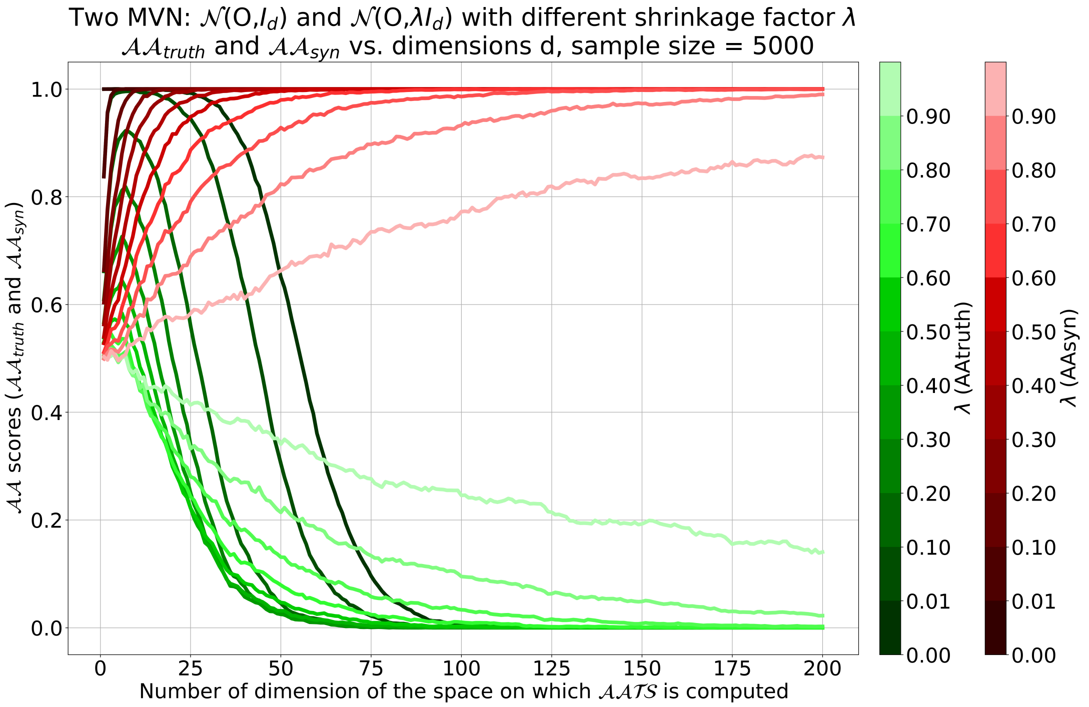


Figure S8: **Effect of shrinkage on $\mathcal{AATS}$.** 5000 samples generated from multivariate normal distributions $\mathcal{N}(0, \mathbb{I}_D)$ (dataset $T$) and $\mathcal{N}(0, \lambda\mathbb{I}_D)$ (dataset $S$), varying the amount of shrinkage $\lambda$ from 0 (extreme shrinkage) to 0.9 (weaker shrinkage)). $\mathcal{AA}_{truth}$ is represented in gradient of green, $\mathcal{AA}_{syn}$ is represented in gradient of red. The color gradient represents the different values of the shrinkage factor. The x-axis is the number of dimensions of the space on which $\mathcal{AATS}$ is computed. On the y-axis are the $\mathcal{AA}$ scores ($\mathcal{AA}_{truth}$ and $\mathcal{AA}_{syn}$).

### S4 Implementation details of PCA-WGAN

Motivated by the experiments done in Section 3.1, in which we observed that the reconstruction error is low enough and reasonably similar when more than 50% of the PCA axes are kept ( **FIG.2**), we decided not to keep all the PCs due to parameter size complexity. We thus designed a GAN that takes as input only 90% of the components, *i.e.* 4507 PCs.

The fully-connected architecture of the generator is the following :

- input layer : 5408 neurons
- 1st hidden layer: 5150 neurons
- BatchNorm1D
- LeakyReLU with negative slope of 0.01
- 2nd hidden layer: 4828 neurons
- BatchNorm1D
- LeakyReLU with negative slope of 0.01
- last layer: 4507 neurons

The fully-connected architecture of the critic is the following :

- input layer : 4507 neurons
- 1st hidden layer: 2253 neurons
- LeakyReLU with negative slope of 0.01
- 2nd hidden layer: 1502 neurons
- LeakyReLU with negative slope of 0.01
- last layer: 1 neuron

For both neural networks the optimizer is RMSprop with weight decay=1e-4. The learning rate for the generator (resp. critic) is 0.0001 (resp. 0.0008). The PCA-WGAN was trained for 1300 epochs. The batch size was set to 32. The noise fed to the generator is a multivariate normal vector with mean 0 and unit variance.

The generator takes input noise of at least the same size as the number of variables of the target distribution and outputs vectors of dimension the number of PCs. The intuition behind this is that it is rather tedious to fill a volume (hypothetical target distribution) with a curve (over-simplified input noise) (see Peano's space-filling curve). The critic takes as input a vector of 4507 dimensions, which is either generated ($\tilde{z}$), or the projection of a real sample onto the eigensubspace ($z$) and outputs a "realness" score.

The critic minimizes

$$\mathbb{E}_{\tilde{z}\sim\mathbb{P}_{generated}}[C(\tilde{z})] - \mathbb{E}_{z\sim\mathbb{P}_{real}}[C(z)] + \lambda\mathbb{E}_{\hat{z}\sim\mathbb{P}_{\hat{z}}}[(\|\nabla_{\hat{z}}C(\hat{z})\| - 1)^2]$$
$$\text{s.t. } \hat{z} = t\tilde{z} + (1-t)z \text{ and } t \sim \mathbb{U}[0,1],$$

while the generator maximizes

$$\mathbb{E}_{\tilde{z}\sim\mathbb{P}_{generated}}[C(\tilde{z})]$$

### S5 Implementation details of Glocal-PCA-WGAN

The SNP matrix was evenly split (along the SNPs) into $K$ parts. Since this approach makes the number of parameters of the model scale with the number of blocks, $K$ was chosen to be small ($= 3$). As before, remarking that the reconstruction error is low enough and reasonably similar around 50% of kept PCA axes (**FIG. 2**), we decided not to keep all the PCs due to parameter size complexity. We thus retained 2000 principal components (i.e. 40% of PCs) per block amounting to ∼90% of the explained variance for each.

The generator takes as input noise of dimension 6000 and output vectors of dimension 6000. In this model, there are as many local critics as there are blocks, plus a global critic. Each local critic aims to capture the local information of its assigned block while the global critic should capture the relations between the blocks. The global critic takes as input either the concatenated projections (1,2,3) or the generated data $(\hat{1}, \hat{2}, \hat{3})$ and outputs a "realness" score. Each local critic takes as input the projection of a block onto its principal subspace truncated to 2000 dimensions, or the corresponding block of generated samples, and also outputs a "realness" score. Each local critic $i$ minimizes the Wasserstein loss restricted to its block. The global critic minimizes the Wasserstein loss restricted over the concatenated samples. The generator maximizes the outputs of the local critics on their assigned blocks (by splitting the generated PC scores) and maximizes the output of the global critic.

Glocal-PCA-WGAN was trained with the same set of hyperparameter values as for PCA-WGAN with a number of epochs set to 1100. For the generator, the number of neurons remains constant throughout the layers.

### S6 Light PCA-WGAN with NN architecture

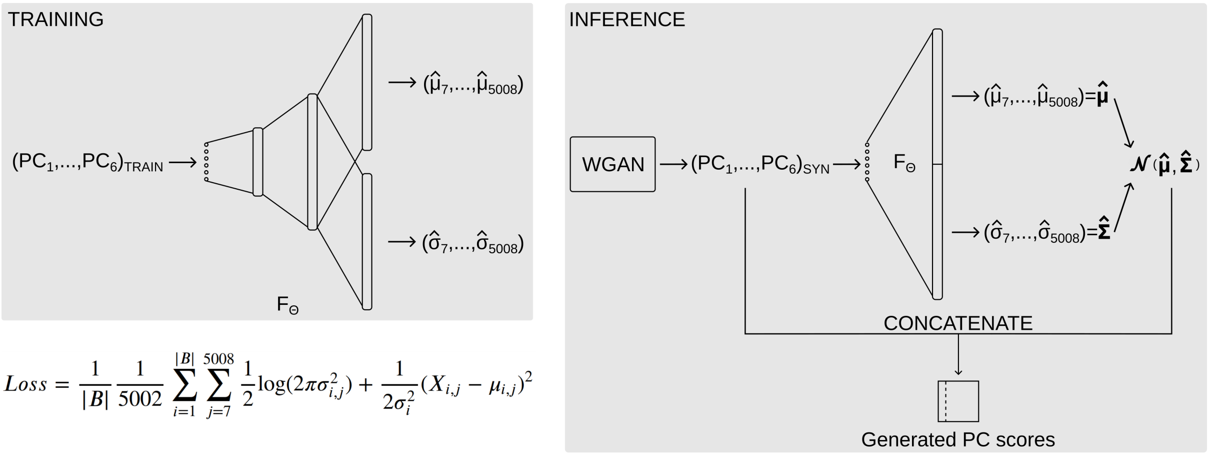


Figure S9: **Light PCA-WGAN with NN.** During training, the neural network was fed with the first six PC scores of the real data and tried to predict the means and variances of the remaining 5002 PC scores. This network was optimized through minimization of Gaussian negative log likelihood loss. During inference, we sampled the Light PCA-WGAN to generate samples with six PC scores, ran them through the trained network, yielding the corresponding means and variances for the 5002 PCs. Finally, we sampled a (diagonal) Gaussian with these generated parameters and concatenated the high and low variance PC scores.

The architecture of $F_\Theta$ (**FIG.S9**) is the following :

- input layer : 6 neurons
- 1st hidden layer: 64 neurons
- BatchNorm1D
- LeakyReLU with negative slope of 0.01
- Repeated sequence of 8 layers, each containing 64 neurons, BatchNorm1D and LeakyReLU with negative slope of 0.01
- last layer: 5002 neurons
- vector of 5002 parameters

The last layer is repeated twice, 5002 neurons for the means and 5002 neurons for the variances. The last layer corresponding to the means is initialized to zero values as PCA is centering the data to the origin beforehand. After running through the last layer, the output is multiplied by a vector of positive, learnable parameters that act as a scaling vector. Adam optimizer is used with a learning rate scheduler, namely, cosine annealing with warmup restart. The parameters for the scheduler are $T_0 = 15, T_{mult} = 1, \eta_{min} = 3e - 12$ and warmup learning rate set to 0.03. A warmup scheduler was used from library "pytorch_warmup", namely, LinearWarmup with warmup period set to 15, i.e., it reaches a learning rate of 0.03 at epoch 15. This network was trained for 100 epochs with GaussianNLL loss from pytorch and a batch size of 5008, i.e., full batch. The training time takes about 30 seconds.

### S7 Light PCA-WGAN with independent Gaussian

We fitted a multivariate normal distribution (MVN) to the real training data by setting its parameters to the means and variances of the remaining 5002 PC scores *i.e.*, PCs with less prominent modal structure. This multivariate normal distribution has diagonal covariance. In this approach which we call Light PCA-WGAN with MVN, we sample the Light PCA-WGAN part and the MVN independently of each other, then concatenate the high and low variance part of a sample, and finally inverse transform it with PCA.
Light PCA-WGAN with MVN or NN provided similar results on the population genetics summary statistics so we only presented results from the latter approach. More details about the relevance of this variant are in AppendixS8 .

### S8 Dependencies between high and low variance PC scores

The mutual information between high and low variance PC scores was computed to assess the dependencies between these variables (**FIG. S10**). The MI for multivariate gaussian with diagonal covariance was used as a null model to evaluate the significance of MI values. The MI for real data PC scores was higher than the null model, showing that these high and low variance variables were not independent. Moreover, the NN (**FIG. S9**) can capture these dependencies. This analysis highlighted the relevance of the Light PCA-WGAN with NN variant.

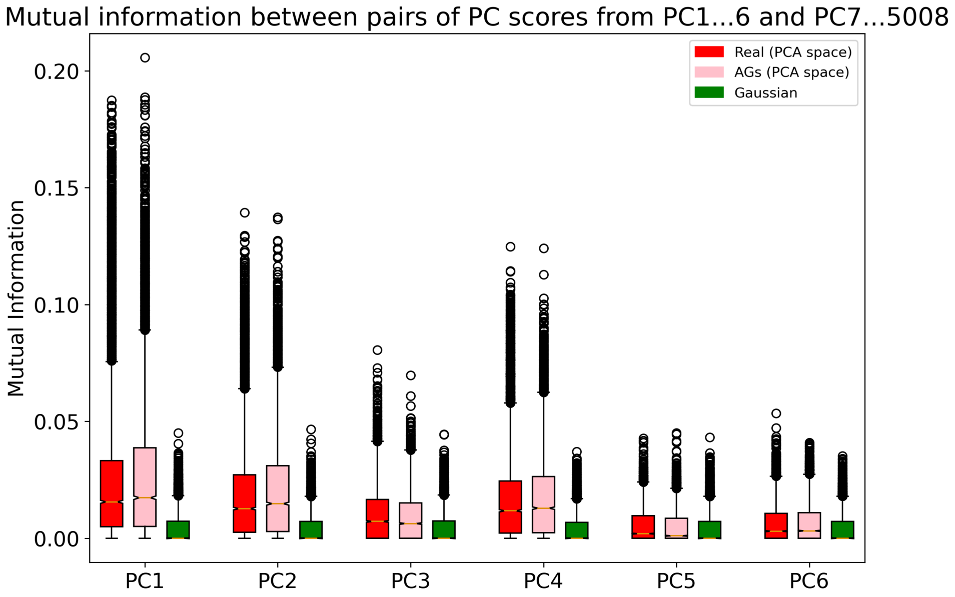


Figure S10: **Mutual information.** Mutual Information (MI) of pairs of high and low variance PC scores for the real data (red), the generated PC scores from Light PCA-WGAN with NN (pink), and a baseline gaussian distribution (green). For a PC score of rank $i \in 1, ..., 6$, we computed its MI with every PC scores of rank $i \in 7, ..., 5008$.

### S9 Parameter size comparison

For PCA-WGAN, the model has parameter size equal to (**FIG. S11**):

- 73M for 10K SNPs
- 73M for 65K SNPs
- 73M for 1M SNPs

For CRBM (Yelmen et al., 2023), the model has parameter size equal to :

- 10M for 10K SNPs
- 130M for 65K SNPs
- 2.08B for 1M SNPs

For convWGAN (Yelmen et al., 2023), the model has parameter size equal to :

- 9.5M for 10K SNPs
- 16.6M for 65K SNPs
- 39.3M for 1M SNPs

For a theoretical WGAN with fully connected architecture (FC-WGAN), the model would have a parameter size equal to :

- 366.7M for 10K SNPs
- 15.7B for 65K SNPs

- 4T for 1M SNPs

For a Glocal-PCA-WGAN, the model have a parameter size equal to :

- 140M keeping 40% of PCs for 65K SNPs
- 3.8B keeping 40% of PCs and 140M keeping 8% of PCs for 1M SNPs and $K = 16$ so that the SNP blocks are of sizes 65535.

We would like to stress out that for a sequence of 1M SNPs, the implementation of the models and their training was not carried out, and is thus purely theoretical. Extrapolating the architecture to higher sequence length does not guarantee a successful convergence during training. Moreover, performing dimension reduction with fixed sample size and increasing the number of features will likely lead to a break down of PCA.

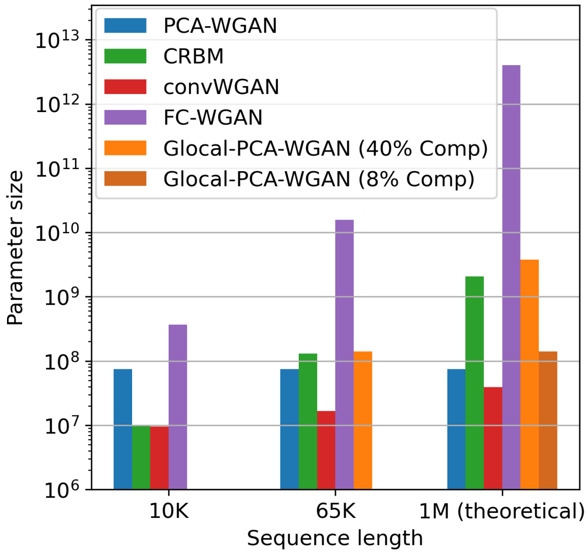


Figure S11: **Complexity & scalability.** Parameter size of the models as a function of sequence length. All models for sequence length of 1M SNPs are theoretical as they were not tested. The CRBM (Yelmen et al., 2023) consists in 13 RBMs having 10M parameters each. The training time of the whole set of RBMs depends on the number of available GPUs. As they can be trained in parallel or in sequence, the complexity is more a matter of time rather than parameter size.

### S10 Analysis of k-mers on particular continental ancestry groups.

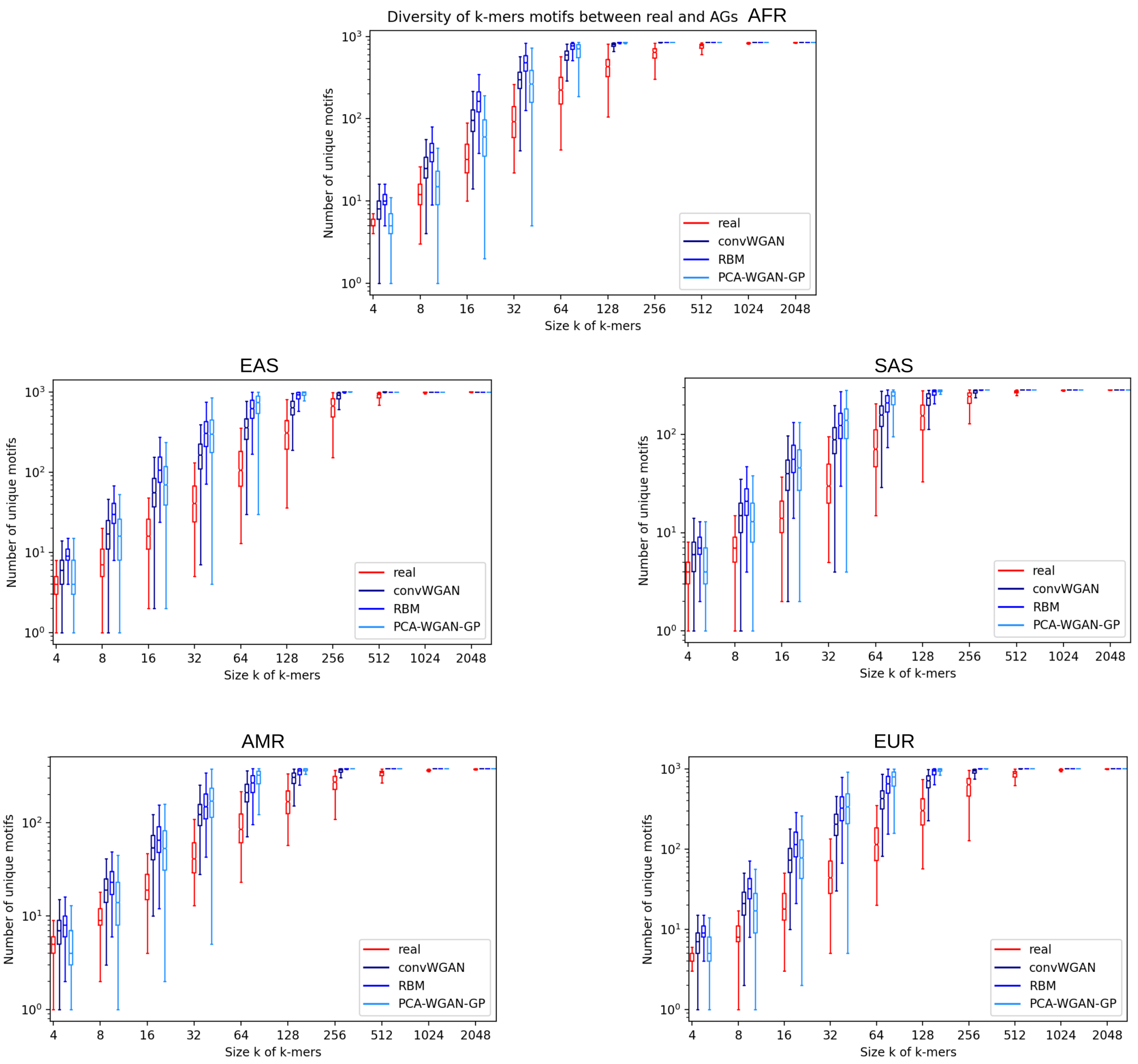


Figure S12: **Diversity by continental ancestry group.**

### S11 Population genetics summary statistics

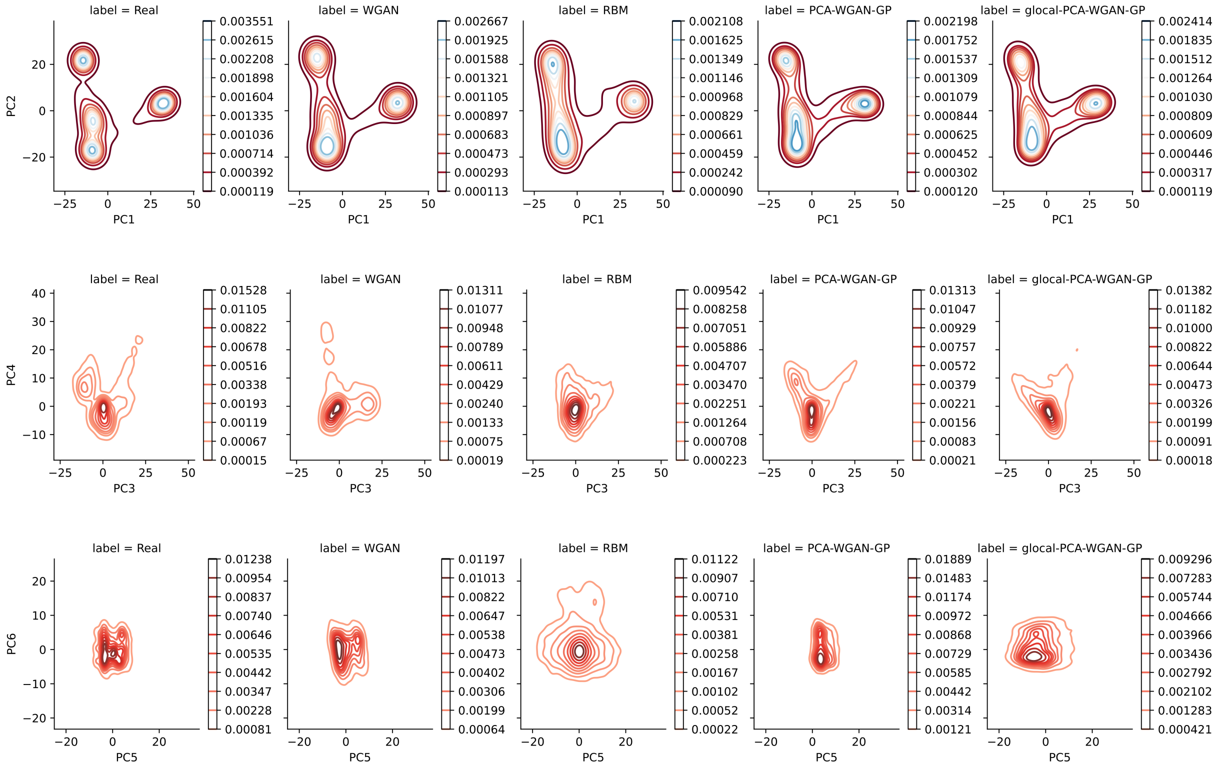


Figure S13: **Density plot of combined real and artificial genome datasets for the first six PCs.** Density increases from red to blue.
